# SARS-CoV-2 Spike Protein Mutations and Escape from Antibodies: a Computational Model of Epitope Loss in Variants of Concern

**DOI:** 10.1101/2021.07.12.452002

**Authors:** Alice Triveri, Stefano A. Serapian, Filippo Marchetti, Filippo Doria, Silvia Pavoni, Fabrizio Cinquini, Elisabetta Moroni, Andrea Rasola, Francesco Frigerio, Giorgio Colombo

## Abstract

The SARS-CoV-2 spike (S) protein is exposed on the viral surface and is the first point of contact between the virus and the host. For these reasons it represents the prime target for Covid-19 vaccines. In recent months, variants of this protein have started to emerge. Their ability to reduce or evade recognition by S-targeting antibodies poses a threat to immunological treatments and raises concerns for their consequences on vaccine efficacy.

To develop a model able to predict the potential impact of S-protein mutations on antibody binding sites, we performed unbiased multi-microsecond molecular dynamics of several glycosylated S-protein variants and applied a straightforward structure-dynamics-energy based strategy to predict potential changes in immunogenic regions on each variant. We recover known epitopes on the reference D614G sequence. By comparing our results, obtained on isolated S-proteins in solution, to recently published data on antibody binding and reactivity in new S variants, we directly show that modifications in the S-protein consistently translate into the loss of potentially immunoreactive regions. Our findings can thus be qualitatively reconnected to the experimentally characterized decreased ability of some of the Abs elicited against the dominant S-sequence to recognize variants. While based on the study of SARS-CoV-2 Spike variants, our computational epitope-prediction strategy is portable and could be applied to study immunoreactivity in mutants of proteins of interest whose structures have been characterized, helping the development/selection of vaccines and antibodies able to control emerging variants.

## Introduction

Protein sequences evolve as a result of selective pressure to optimize function, create improved phenotypes, and introduce new advantageous traits. In pathogens like bacteria and viruses, sequences evolve via modifications such as point mutations, recombination and deletions/insertions to induce higher infectivity, more efficient replication, and ultimately escape from the host immune systems ^[1]^.

The SARS-CoV-2 virus, the etiological agent of COVID-19, is no exception to these general rules. The spread of the virus to more than 200 million people worldwide, combined with the pressure determined by the reactions of immunocompetent populations, led to the emergence of “variants of concern”. In this context, attention has been focused on the SARS-CoV-2 spike protein (S protein), the large, heavily glycosylated class I trimeric fusion protein which mediates host cell recognition, binding and entry. Because it represents the first point of contact with the host, and given its crucial role in viral pathogenesis ^[1e, 1f, 2]^, the S protein has been the basis for the design of currently used vaccines effective at reducing viral spread, hospitalization and mortality rates ^[3]^.

While for almost one year the only notable mutation in S has been the D614G (Asp^614^ →Gly), which increases affinity for the cell receptor ACE2 and has immediately become dominant, novel S protein variants reported of late may pose new potential challenges for efficacy of vaccination, antibody-based therapies and viral diffusion control. Three notable examples of such evolved S proteins, which correspond to major circulating variants, are B.1.1.7 (the so-called UK or alpha variant), 501Y.V2/B.1.351 (the South African or beta variant), and B.1.1.28 (P.1, the Brazilian or gamma variant). All such sequences contain various mutations due to nonsynonymous nucleotide changes in the RBD domain, including E484K, N501Y, and/or K417N ^[2c]^. In B.1.1.7 and B.1.351, deletions are also present in the N-terminal Domains (**Figure 1**).

**Figure 1.**
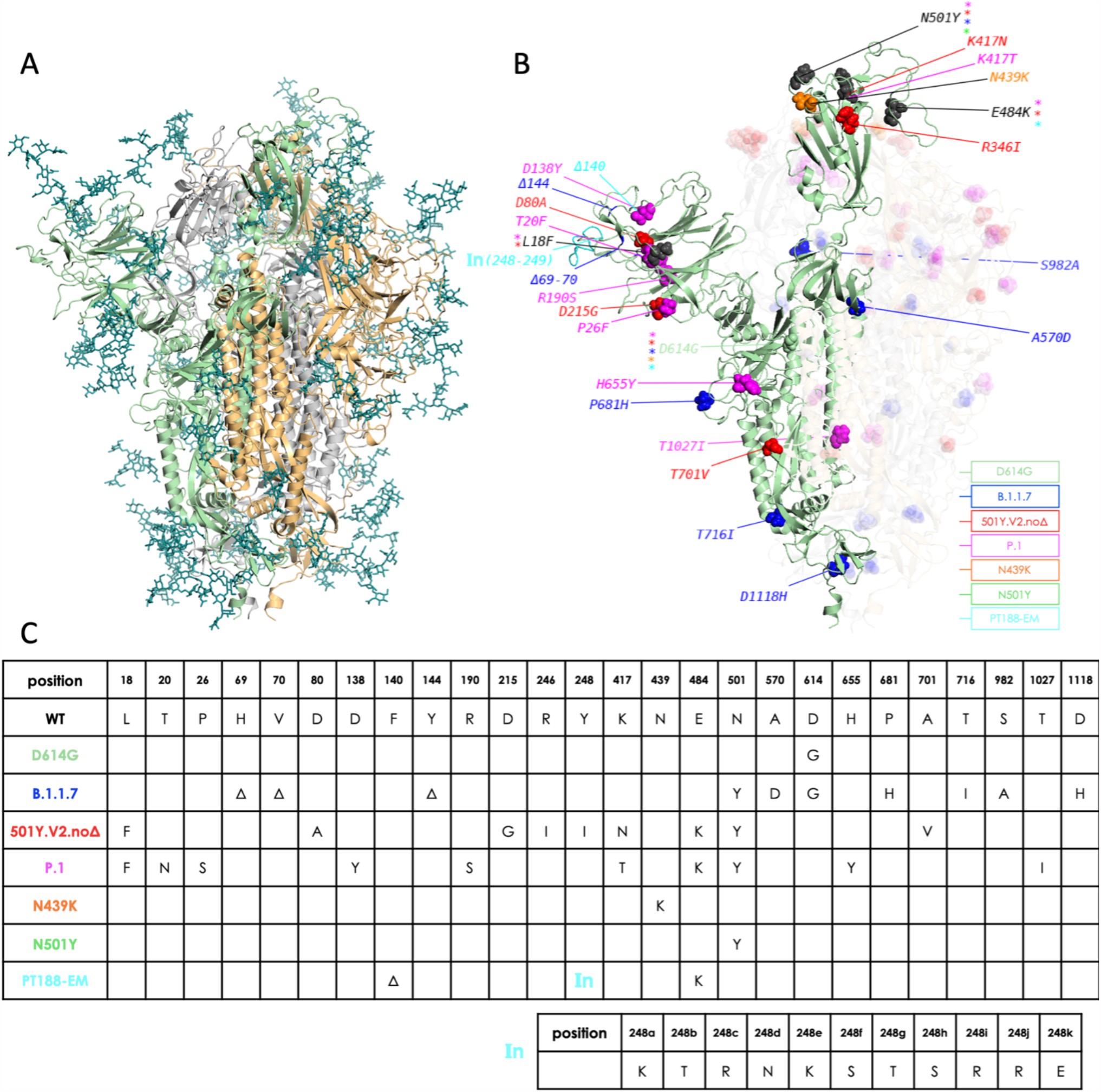
Overview of simulated variants (definitions in main text). **A)** The full-length, fully glycosylated trimeric structure corresponding to pdb code 6VSB. Protomer A (RBD “up”): secondary structure in green; protomers B and C (RBD “down”): grey and sand, respectively. Glycans’ C, N, and O atoms rendered as teal sticks. **B)** Positions and nature of mutations highlighted on protomer A of different variants. Mutant residues’ heavy atoms are rendered as spheres; a different color is assigned to each variant, as indicated in the legend. Mutations common to more than one variant are rendered and/or labeled in black, with colored asterisks denoting variants carrying the mutation. The insertion in the PT188-EM variant (cyan) is denoted by “**In**(248-249)”. Protomers B and C are also shown with their respective mutations, but rendered with increased transparency for clarity; glycans are omitted**; C)** Synopsis of mutations on the different variants simulated in this work, including the 11-residue insertion in the PT188-EM variant.

Several studies showed how some of these circulating variants may have reduced sensitivity to neutralizing antibodies targeting the RBD or to the NTD ^[2c, 4]^. In this context, polyclonal antibodies contained in Convalescent Plasma (CP) from individuals infected with the D614G-containing SARS-CoV-2, showed reduced potency in neutralizing 501Y.V2/B.1.351 virus isolates ^[5]^. Furthermore, antibodies elicited after vaccine treatment showed reduced neutralization of pseudoviruses bearing the mutations of the P.1 and 501Y.V2/B.1.351 variants ^[6]^. The same was observed for pseudoviruses with variations in S mimicking those of the B.1.1.7 lineage ^[6-7]^. Yet, fortunately, it was shown that vaccine-generated antibody titers were sufficient to neutralize B.1.1.7 in sera from 40 BNT162b2-vaccinated individuals ^[8]^. In this context, it is encouraging to note that new studies are reporting high levels of efficacy against severe forms of Covid-19 also in countries where these variants have become dominant ^[9]^.

A crucial question for understanding the impact of S-protein evolution on the development of monoclonal antibody (mAb)-based and vaccine-based therapies, is whether we can develop a simple model to rationalize, and eventually predict, the effect of variations on the structural properties of S that ultimately underpin antibody recognition. Fundamentally, comparison across S-proteins mutants can help us understand the molecular basis of the protein’s evolvability, furthering our grasp of the relationships between sequence, structure and (immuno)recognition. From the practical point of view, this knowledge could in principle be harnessed to design and engineer improved S-based antigens or multicomponent domain/peptide combinations, focusing for instance on those antibody binding regions, known as epitopes, that are predicted to be conserved in multiple variants.

Here, we apply a straightforward structure-dynamics-energy strategy to predict potentially immunogenic regions in representative 3D conformations of several variants of the full-length glycosylated trimeric S protein (**Figure 1**).

The selected S proteins represent some of the major variants of concern circulating at the time of setting up simulation. In this respect, the African variant we simulate, which is named 501Y.V2.noΔ,, corresponds to the S lineage originally discovered in South Africa in late November 2020 by Tegally *et al*.^[10]^. This S variant features the additional mutations L18F (in common with P.1) and R246I but does not feature the Δ241-243 deletion, whose existence was still debated when the authors released their study in January 2021 ^[10]^. This variant has subsequently been referred to in several papers as B.1.351^[5]^. The list of studied proteins is further enriched by a laboratory-evolved escape S-variant, obtained by Rappuoli and coworkers by co-incubating the SARS-CoV-2 virus with a highly neutralizing plasma from a COVID-19 convalescent patient. Interestingly, after several passages this strategy generated a variant completely resistant to plasma neutralization. This “artificial” variant is labeled here as the PT188-EM variant ^[5b]^.

Conformations are extracted from independent atomistic molecular dynamics (MD) simulations totaling 4 µs for each mutant. Our approach to the detection of epitopes on S, i.e., its antibody-binding protein regions, is based on the concept that such sites should continuously evolve to escape immune recognition by the host without impairing the native protein structure required for viral function and survival. We previously showed—and experimentally confirmed— that these regions coincide with substructures that are not involved in major stabilizing *intramolecular* interactions with core protein residues that are important for its folding into a functional 3D structure ^[11]^. In other words, Ab-interacting regions show minimal energetic coupling with the rest of the protein, which in turn should favor accumulation of escape mutations while preserving the antigen’s 3D structure. Furthermore, minimal intramolecular coupling provides epitopes with greater conformational freedom to adapt to and be recognized by a binding partner. Actual binding to an external partner such as an Ab is expected to occur if favorable intermolecular interactions determine a lower free energy for the bound state than for the unbound state^[11-12]^.

These concepts are analyzable by the MLCE approach ^[11, 13]^ (see also methods). Starting from the characterization of the energy of pairwise interactions between all aminoacids and monosaccharides, and filtering the resulting interaction map with structural information extracted from the same protein’s inter-residue contact map, MLCE identifies groups of spatially contiguous residues with poor energetic coupling to the rest of the protein as potential immunogenic regions. At the same time, groups of residues with high energetic coupling are identified as stabilization centers.

Upon comparing our results to recently reported characterization of Ab binding and reactivity, the analysis we report consistently shows that mutations, deletions, and/or insertions in S variants determine a reorganization of internal interactions leading to the loss of potentially immunoreactive regions on the surface. Encouragingly, these findings can be qualitatively reconnected to the decreased ability of some of the Abs elicited against the dominant S-sequence to recognize variants.

## Results

To characterize the effects of mutations, deletions, and insertions on the definition of potential Ab-binding substructures in S variants, we apply a combination of the Energy Decomposition (ED) and MLCE (Matrix of Low Coupling Energies) methods ^[11, 13-14]^ to representative structures extracted from long timescale MD simulations of the S protein variants reported in **Figure 1**.

Briefly, we first run 4 independent 1 µs long all-atom MD simulations of each variant of the full-length fully glycosylated S protein in solution (**Figure 1**) (each built from PDB ID: 6VSB ^[3a]^). Next, for each variant, we concatenate individual trajectories into one a single 4 µs metatrajectory. Cluster analysis on each variant’s metatrajectory is then conducted to identify the 3 most representative conformations. These are then used to compute nonbonded pairwise potential energy terms (van der Waals, electrostatic interactions, solvent effects) obtaining, for a given variant with *N* aminoacid and monosaccharide residues, a symmetric *N* × *N* inter-residue interaction matrix. The three matrices extracted from a variant’s trajectory are then weighted and averaged to yield an average nonbonded interaction matrix, *M*_*ij*_. Upon eigenvalue decomposition of *M*_*ij*_, eigenvectors associated with the most negative eigenvalues can help build a simplified version of *M*_*ij*_ that only highlights series of residues with high- and low-intensity couplings. The former represent residues acting as folding hotspots and responsible stabilizing the protein’s 3D structure; the latter represent residue pairs with weak energetic coupling to the rest of the protein, whose mutation is expected not to impact S’ structural functionality. In this framework, once information contained in the simplified energy map is combined with information contained in the protein’s residue-residue contact map, it permits to ‘filter out’ clusters of residues whose energetic coupling to the rest of the structure is weak *and* that are spatially contiguous. Such localized networks of low-intensity couplings, located in proximity of the protein surface represent potential interaction Ab-interaction regions, or epitopes. This approach has been previously experimentally validated in a number of applications^[12a, 12b, 15]^.

The reference S structure we use here is the dominant D614G variant. We analyze the results of epitope predictions we obtain on *isolated* S variants by comparing them against selected Spike-antibody complexes. To this end, we collected X-ray or Cryo-EM structural data of complexes between S and various Abs, reported in **Table 1 and Table S1**. Epitopes in experimental structures are defined as the sets of S protein residues within 5Å of any Ab residue **(see Supporting Information Table S2)**. The experimental epitopes thus derived are used as the reference against which to compare epitopes predicted *in silico*.

**Table 1.**
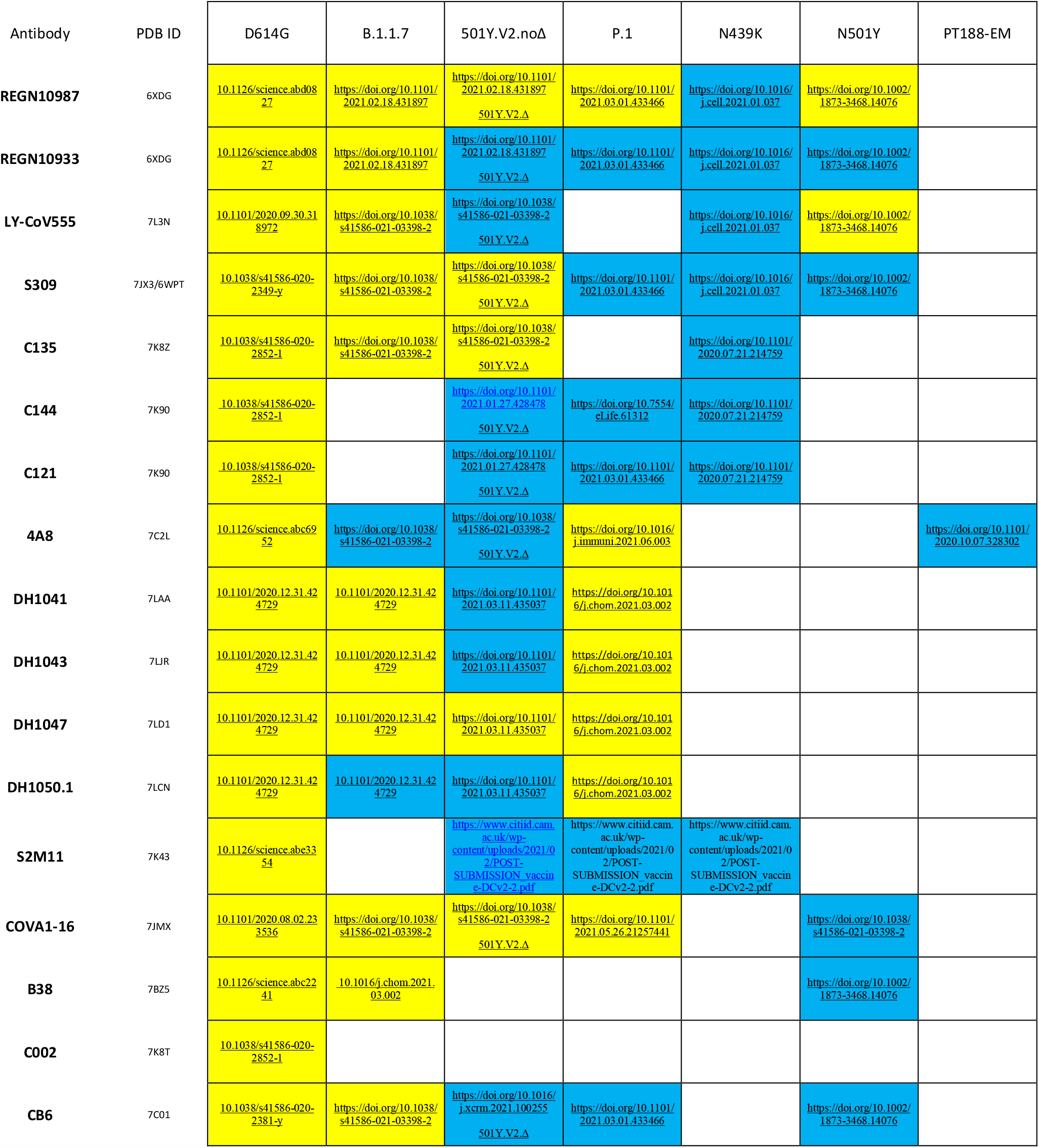
PDB IDs of the S-Ab complexes used to compare epitope predictions. For each Ab considered in this work (leftmost column), we report: PDB IDs of S-Ab Cryo-EM complexes used as experimental reference for our MLCE epitope predictions; and, where available, experimental studies reporting either that Ab’s gain (yellow) or loss/absence of activity (blue) towards a particular variant. White cells indicate that experimental data is unavailable. * denotes experimental studies carried out on the 501Y.V2.noΔ S variant but with the Δ241-243 deletion.

Figure 2 schematically reports the sequences of the RBD and N-terminal domain for each variant studied (sequences on the Y-axis, variant on the X-axis). The different colors point out the residues of a certain variant that are predicted to be part of an epitope for a certain characterized antibody. Importantly, for the reference variant predicted epitopes largely overlap with experimentally identified regions. In particular, epitopes are correctly predicted for Abs targeting both the Receptor Binding Domain (RBD) and the N-terminal domain of the protein (**Table S2, Figure 2**).

In the remaining variants of concern, a diverse landscape of epitopes emerges. A number of residues/regions that are predicted immunogenic in the reference S-protein disappear in the variants. Overall, this is observed for all the Abs considered.

**Figure 2.**
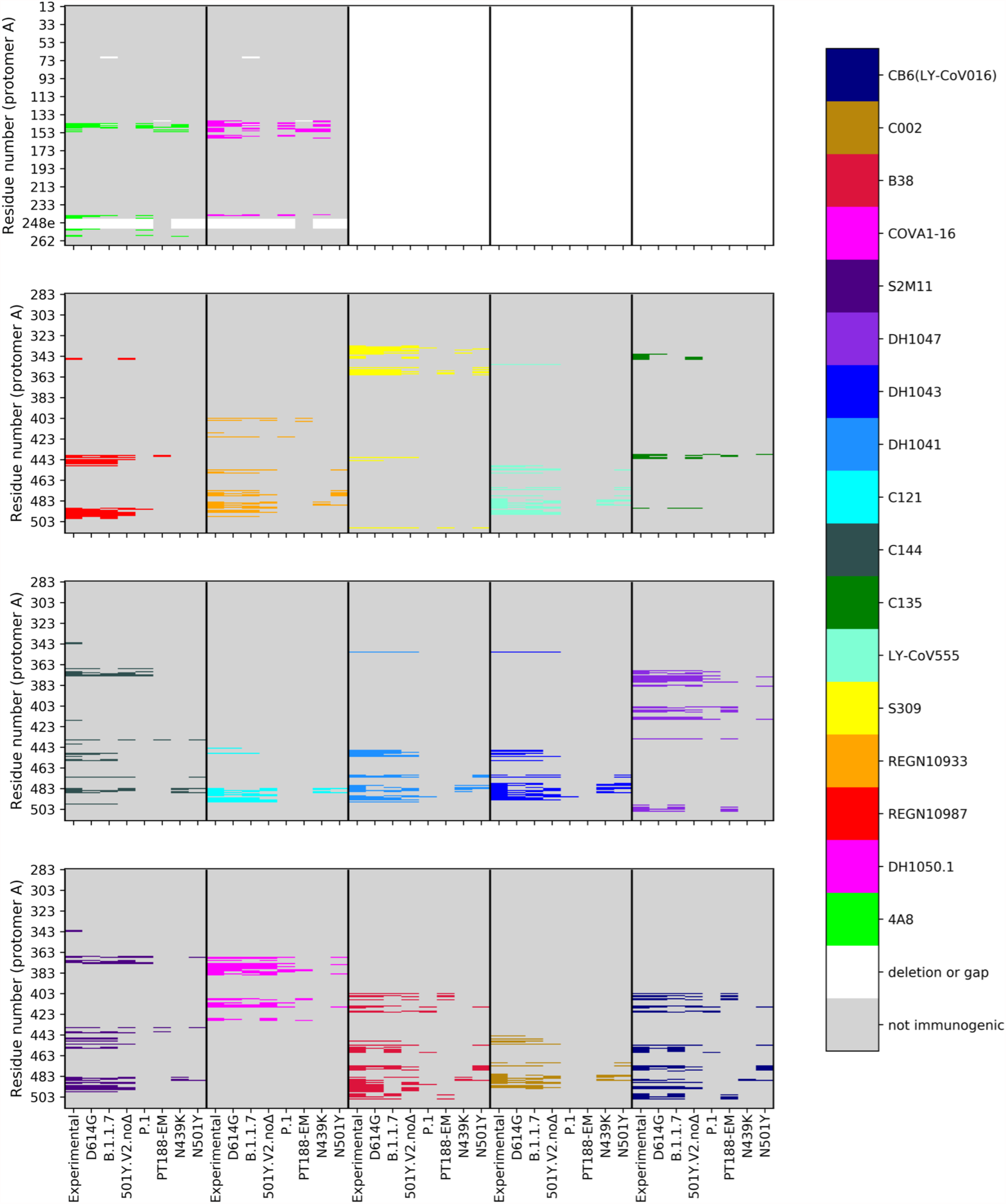
Mapping epitopes on each variant. Epitope mapping on S protomer A for the 2 NTD-targeting antibodies (two top left panels; *cf*. numbering on *y-*axis) and the 15 RBD-targeting antibodies (bottom three rows) considered in this study. In each panel, using a distinct color for each antibody (right palette), epitopic residues detected in the Cryo-EM or crystal structures (labeled “Experimental” on each panel’s *x-*axis) are compared to epitopes predicted *in silico* on each of the seven variants considered. Non-immunogenic residues are shown in gray; gaps/insertions in white.

In this framework, after running an epitope prediction on each variant we monitor epitope conservation across variants through a *conservation ratio*: the number of residues in each predicted epitope for a given variant is divided by the number of residues in the corresponding experimental epitope in the reference S structure, which is defined based on the 5 Å threshold from its respective Ab, as discussed above. We define *epitope loss* when the conservation ratio is lower than 0.5; otherwise the epitope is considered to be conserved. In **Table 2** and **Figure 3** we report such conservation ratios for each D614G S epitope on each simulated variant, and confront them with available experimental data (at the time of writing) on the variant’s reactivity towards the Ab that would be expected to bind to that particular epitope. Each cell in the table is color-coded according to the experimentally measured activity of the corresponding Ab on one of the given variants. If the Ab remains active, the cell is yellow. If the Ab has lost activity against that variant, the cell is blue. If experimental data is unavailable for a particular Ab on a particular variant, the cell is white.

**Table 2.**
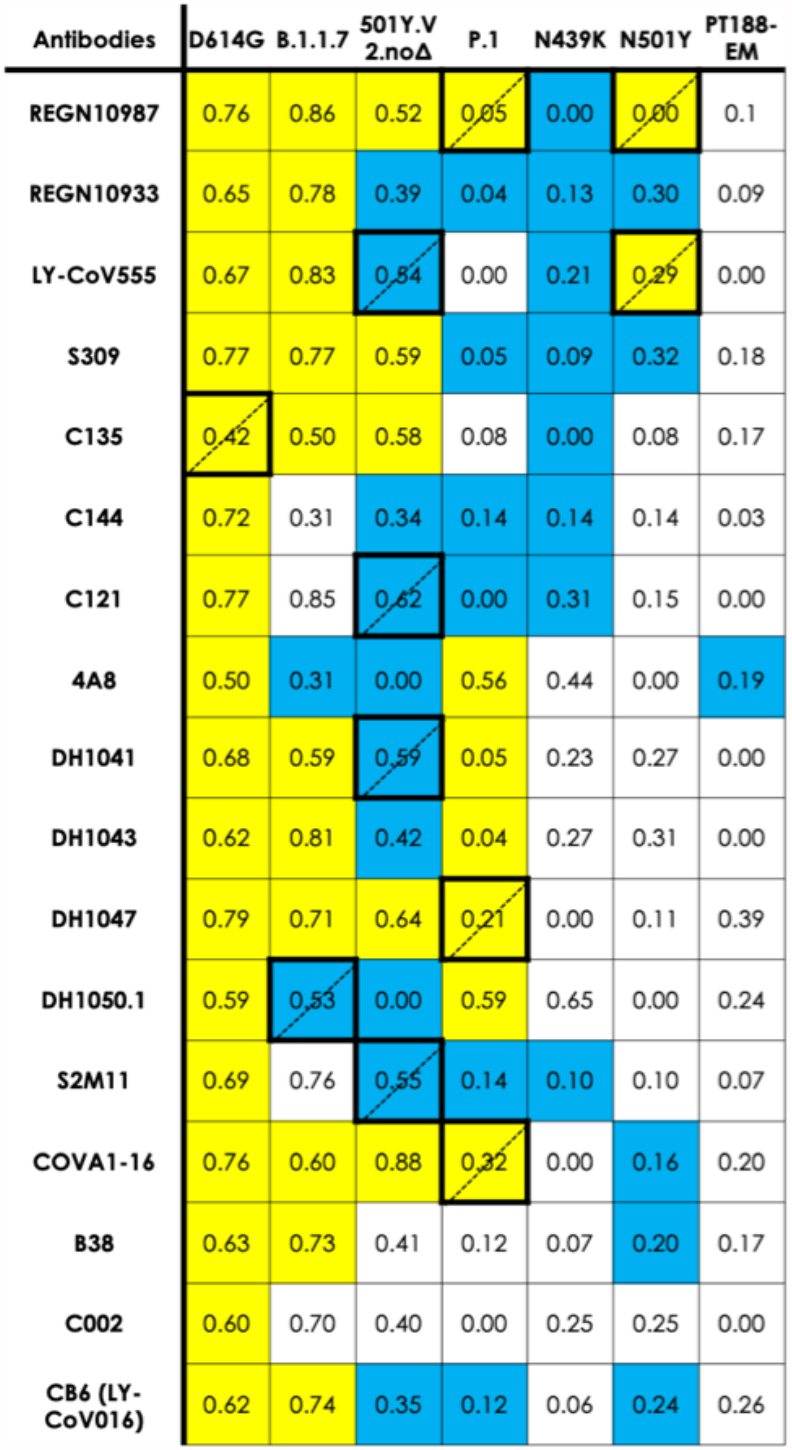
Epitope predictions on each variant and epitope conservation ratio. Each cell reports an *epitope conservation ratio* for each S variant-Ab combination, relating *in silico* predictions to experimental epitopes from experimental Cryo-EM and/or crystal structures. Conservation ratios lower than 0.5 indicate epitope loss; otherwise an epitope is considered to be conserved. Each cell in the table is color-coded according to the experimentally measured activity of the corresponding Ab on the respective variant. If the Ab remains active, the cell is yellow. If the Ab has lost activity against that variant, the cell is blue. If experimental data is unavailable for a particular Ab on a particular variant, the cell is white. Disagreement between predictions and experiment (i.e., blue and conservation ratio >0.5 or yellow and conservation ratio <0.5) is indicated by thick borders and dotted-line diagonal.

| Antibodies | D614G | B.1.1.7 | 501Y.V<br>2.noΔ | P.1 | N439K | N501Y | PT188-EM |
| --- | --- | --- | --- | --- | --- | --- | --- |
| REGN10987 | 0.76 | 0.86 | 0.52 | 0.05 | 0.00 | 0.00 | 0.1 |
| REGN10933 | 0.65 | 0.78 | 0.39 | 0.04 | 0.13 | 0.30 | 0.09 |
| LY-CoV555 | 0.67 | 0.83 | 0.84 | 0.00 | 0.21 | 0.29 | 0.00 |
| S309 | 0.77 | 0.77 | 0.59 | 0.05 | 0.09 | 0.32 | 0.18 |
| C135 | 0.42 | 0.50 | 0.58 | 0.08 | 0.00 | 0.08 | 0.17 |
| C144 | 0.72 | 0.31 | 0.34 | 0.14 | 0.14 | 0.14 | 0.03 |
| C121 | 0.77 | 0.85 | 0.62 | 0.00 | 0.31 | 0.15 | 0.00 |
| 4A8 | 0.50 | 0.31 | 0.00 | 0.56 | 0.44 | 0.00 | 0.19 |
| DH1041 | 0.68 | 0.59 | 0.59 | 0.05 | 0.23 | 0.27 | 0.00 |
| DH1043 | 0.62 | 0.81 | 0.42 | 0.04 | 0.27 | 0.31 | 0.00 |
| DH1047 | 0.79 | 0.71 | 0.64 | 0.21 | 0.00 | 0.11 | 0.39 |
| DH1050.1 | 0.59 | 0.53 | 0.00 | 0.59 | 0.65 | 0.00 | 0.24 |
| S2M11 | 0.69 | 0.76 | 0.55 | 0.14 | 0.10 | 0.10 | 0.07 |
| COVA1-16 | 0.76 | 0.60 | 0.88 | 0.32 | 0.00 | 0.16 | 0.20 |
| B38 | 0.63 | 0.73 | 0.41 | 0.12 | 0.07 | 0.20 | 0.17 |
| C002 | 0.60 | 0.70 | 0.40 | 0.00 | 0.25 | 0.25 | 0.00 |
| CB6 (LY-CoV016) | 0.62 | 0.74 | 0.35 | 0.12 | 0.06 | 0.24 | 0.26 |

**Figure 3.**
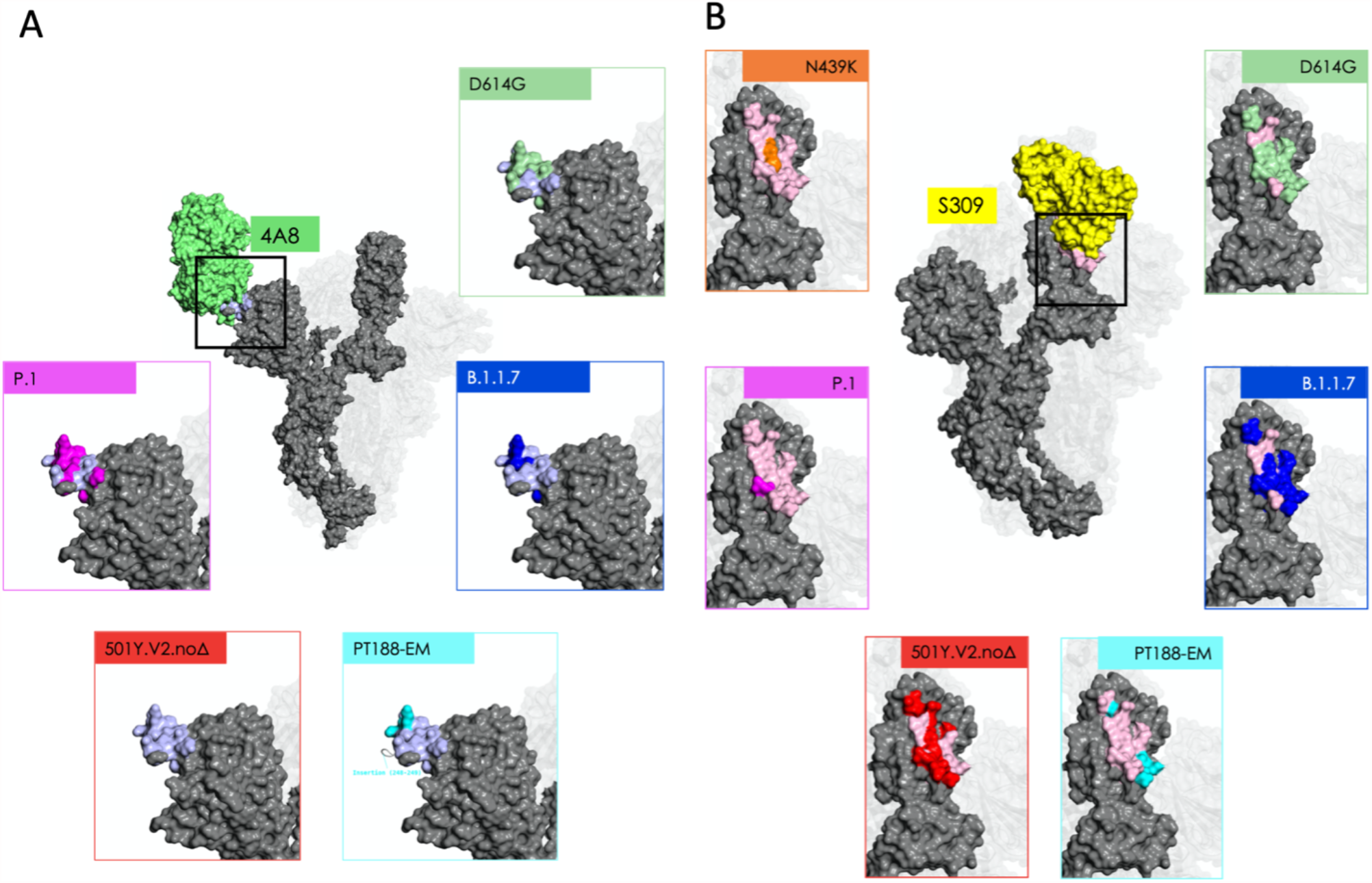
Mutations modify epitope identity. Central images in panels A and B depict the Cryo-EM structure of the antigen-binding fragments of two representative Abs bound to protomer A: 4A8 (panel A; Ab in green; experimental epitope in light blue); and S309 (panel B; Ab in yellow; experimental epitopes in light pink). Insets in each panel contrast the extent of the experimental epitope with epitopic residues predicted by the MLCE method (see main text) for five (panel A) or six (panel B) of the variants considered in this work: these residues are rendered using the same color code used for variants in Figure 1; residues in the experimental epitope not predicted by MLCE are rendered as in the central image (panel A: light blue; panel B: light pink). Other residues on the S protein (not comprised in the experimental epitope) are rendered in gray. Glycans are omitted for clarity; positions of protomers B and C are shown for reference.

Analysis of **Table 2** clearly shows that the vast majority of blue cells, indicative of a loss of Ab reactivity, contain ratios lower than 0.5. This is an important validation of our prediction: whenever a variant’s predicted epitope residues—*i*.*e*., according to MLCE, contiguous residues uncoupled from the S protein core—shrink in number compared to D614G S, it is very likely that experimental data will also confirm that that variant evades Abs binding to the shrunk or lost epitopes. On the other hand, the overwhelming majority of cases for which Abs retain activity against a variant (yellow cells) are also confirmed by our prediction to retain their respective epitopes (conservation ratio > 0.5) with respect to D614G S. Disagreement between our predictions and experiment only occurs in a minority of cases: corresponding cells are marked by thicker borders.

Analysis of B.1.1.7 (UK) and 501Y.V2.noΔ South Africa; late November 2020) immediately shows that a large portion of predicted epitopes in the RBD are conserved compared to the reference D614G. Interestingly, however, we also observe a dramatic drop in the number of NTD residues predicted as epitopes for the 501Y.V2.noΔ. Epitope loss in the NTD, which was deemed to host a super-antigenic hotspot ^[16]^ can help explain the ability for immune evasiveness observed for these two variants. In B.1.1.7, the NTD epitope is largely conserved consistent with the conservation of activity of this Ab against the variant (**Table 2, Figure 2** and **3**).

Importantly, conservation of a dominant part of the epitopes in the RBD still endows the two variants with reactivity against Abs directed to this domain, which may help explain the observed effectiveness of some convalescent plasma treatments and vaccines ^[3c,Yadav, 2021 #19517]^.

Calculations on the Brazilian variant correctly indicate loss of immunoreactivity of several Abs as well as conserved reactivity of Abs 4A8 and S2M11. This variant is the only one for which our predictions of epitopes binding Abs of the DH family generally disagree with experimental data. Finally, it is important to note that the “artificial” PT188-EM variant, evolved in the lab under the pressure of convalescent serum to evade Ab-effects, appears to have lost a very large number of protein epitopes (see **Table 2, Figure 2** and **3**). In particular, the insertion at residues 248 modifies the conformational properties of the region otherwise recognized by Ab 4A8. As a consequence, the epitope to this antibody disappears from the predictions on the PT188-EM variant ^[5b]^. Interestingly, in this case, the carbohydrate motifs coating the protein appear to host most of the uncoupled regions (117 carbohydrate moieties in the PT188-EM variant vs. 90 in the reference S-protein), pointing to a role of the glycan shield in protecting the protein from immune recognition, besides playing a key part in modulating interactions for ACE2 recognition and cell-entry. ^[17]^ (see **Figure 4**).

**Figure 4.**
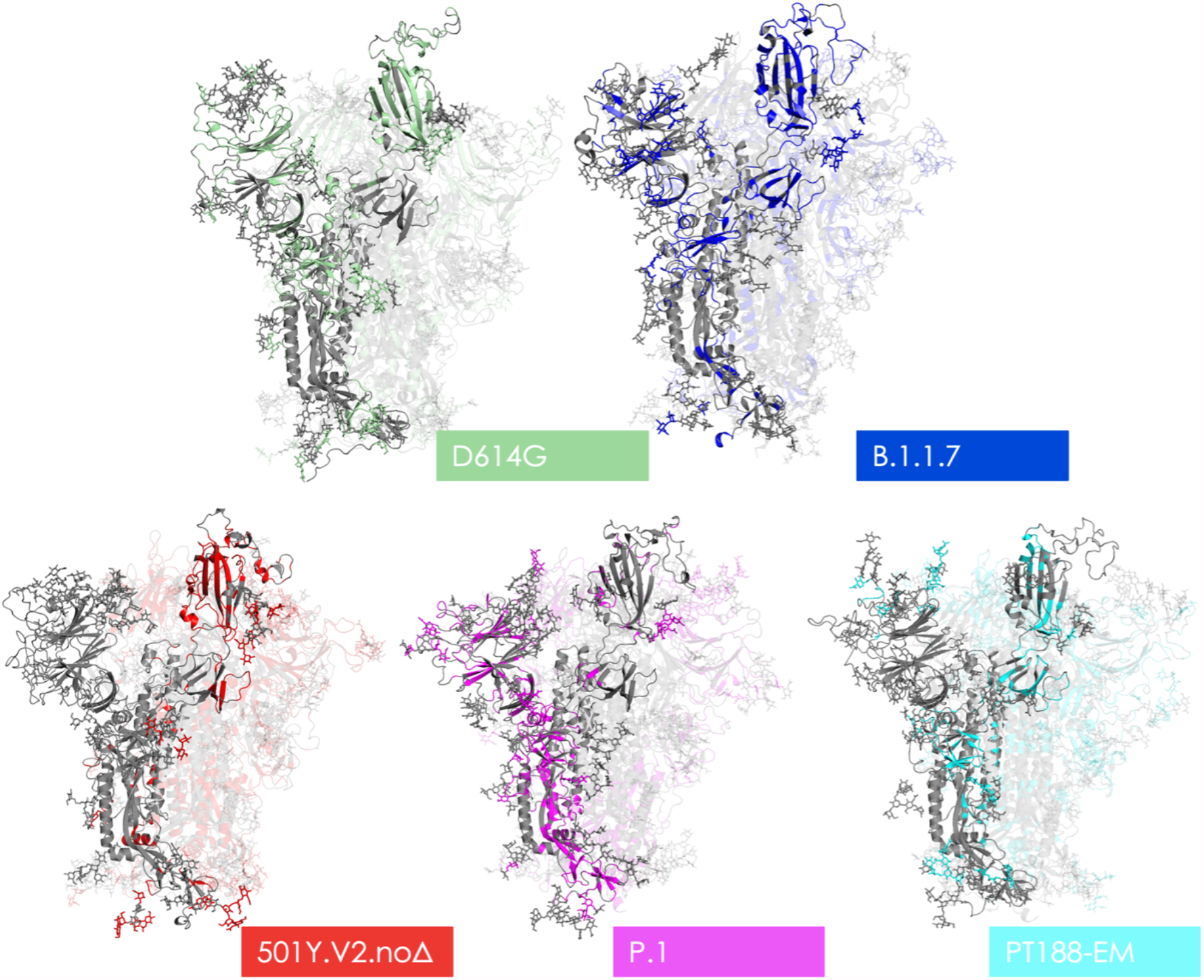
Structural representations of epitopes on different variants. The various structures depicted show the 3D structure of protomer A in gray. Residues rendered in the color assigned to their respective variant in Figure 2 mark the locations of all predicted epitopes; areas in gray represent non-immunogenic regions. Glycan heavy atoms are rendered as sticks.

Importantly, mutants N439K, is correctly predicted as an escape variant from all Abs for which experimental data proved lower efficacy.

## Discussion

In this work, we analyzed full-length models of 7 trimeric glycosylated SARS-CoV-2 S protein variants, derived from the prefusion conformation of the cryo-EM structure 6VSB ^[3a]^, in which the Receptor Binding Domain of chain A (RBD-A) is in an “up” conformation, exposed to interaction with host cell receptors and potential targeting by Abs. The data from our energetic analyses can be aptly integrated in the characterization of the properties of S and other SARS-CoV-2 proteins from long scale simulations, such as those recently presented by Zimmerman et al. coworkers ^[18]^, Casalino et al. ^[17e]^, Spinello et al. ^[19]^, Oliveira et al. ^[20]^, Shoemark et al. ^[21]^, and Wang et al. ^[22]^.

Our MLCE analysis of the full-length trimers correctly identifies a number of epitopes in the RBD that have been previously characterized. RBD is in fact targeted by the largest fraction of neutralizing antibodies. MLCE also identifies regions in the N-terminal domain (NTD), which are known to be targeted by different Abs, some of which potently neutralize SARS-CoV-2 ^[4c]^ and highlights putative immunoreactive substructures at the end of the S2 domain, where sugar-engaging Abs have recently been characterized (see **Table S2, Figure 4**).

Energy-based epitope prediction through the MLCE approach reveals a common theme across variants: the number and surface exposure of potentially immunoreactive regions decrease in S protein mutants compared to the reference D641G. In particular, the number of residues defining the epitope located in the long RBD loop (residues 417-503, recognized by many protective Abs) is much lower in mutants 501Y.V2.noΔ, B1.1.28, and N439K (see **Figures 2, 3, Table S2**). Interestingly, in the case of B.1.1.7, which shows limited evasion, the loop is largely active in terms of immunoreactivity. In contrast, in the evading variant PT188-EM the entire loop disappears from the list of potential Ab-targets.

Experimental characterization of Abs targeting the NTD revealed a site recognized by most Abs, located between the N3 and N5 loops of the domain. This epitope was correctly predicted in our previous work^[15h]^. Specifically, Lys147 and Arg246, known to be important in stabilizing interactions with the complementarity-determining regions of different Abs are correctly predicted as epitope elements.

On the other hand, sequence mutations in SARS-CoV-2 variants lead to the N3 and N5 NTD loops disappearing from the ensemble of Ab-binding substructures. This is observed computationally and is corroborated by recent experimental data by Veesler and coworkers^[23]^. Interestingly, these epitopes largely coincide with the regions where Alanine substitutions reduced affinity for antibodies 4A8, CM17, and CM25 (see ^[23]^) The impact of epitope loss in these regions is also confirmed by the observation that an engineered N3-N5 double mutant and native beta variant ^[10]^ both evade neutralization by mAbs CM25 and 4A8.

Finally, our strategy correctly predicts the loss of most epitopes in the lab-evolved escape variant described by Andreano *et al*. ^[5b]^ (see **Figure 4, Table S2)**.

We propose a model for the study of Ab-reactivity of SARS-CoV-2 S protein variants that integrates sequence and structural information and incorporates dynamics and energetics into the analysis of the variation/loss of epitopes. Mutations in S variants determine the loss of epitopes and as a consequence can confer escape from antibodies. Upon sequence variation, the protein shifts to states characterized by different intramolecular interactions compared to the initial D614G structure; this transition decreases the number of energetically uncoupled substructures available for engaging interactors such as Abs. Unique to this model is the observation that mutations, insertions, and deletions exhibiting different immunoreactivity experimentally are consistently captured by the energy based decomposition of structures extracted from unbiased classical MD simulations of the glycosylated S protein *isolated* in solution, without any input of prior information on Ab-binding propensities. Although qualitative in nature and focused on the study of S variants of concern, our approach is general and immediately portable to other targets to provide physico-chemical information on the determinants of Abs recognition.

Since one of the fundamental goals of structural vaccinology is the identification and design of structures with optimized properties for immunoreactivity, development and validation of computational methods that help identify conserved *vs*. non-conserved epitope regions in different variants independently of whether structures of related protein-antibody complexes are available may hold great potential. In the case we have presented here, one may consider designing chimeras or multicomponent systems (peptide- or domain-based) presenting all (or most of) the conserved sequences that are predicted to be potentially Ab-reactive.

Furthermore, our results suggest that approaches like the one we presented here may be used prospectively as an aid in the analysis and characterization of emerging variants.

Though targeted experiments and design of mutants with tailored reactivities based on MLCE analysis are required to further validate these ideas and precisely define their progression to real-world applicability, our findings provide a new basis to understand how mutations could directly result in escape from immunorecognition.

## Materials and Methods

### Preparation of Spike Protein Variants

Fully glycosylated S protein variants simulated in this work were variously derived from simulations described by Grant *et al*. ^[17b]^ based on the Cryo-EM structure of the WT S protein at PDB entry 6VSB ^[3a]^, wherein one RBD is in the “up” conformation and the other two are “down”. All mutations, including the “reference” D614G, are introduced using the “mutations wizard” in the *PyMOL* molecular modeling package (Schrodinger LLC): rotamers of non-glycine side chains are chosen from the first suggested option for S protomer A, and then, where possible, we have sought to adopt the same rotamers for protomers B and C. Histidine tautomers and disulfide bridges are retained as in our reference simulations. In B.1.1.7 variant S protomers, mutant histidines 681 and 1118 are introduced with protonation at Nε2, and mutant aspartate 570 side chains are left unprotonated. Mutant lysine 484 sidechains (B.1.1.28 variant; E484K variant) are left protonated.

Consistent with our reference simulations, ^[15h, 17b]^ all three protomers are modeled without gaps, from Ala27 in the NTD to Asp1146 just downstream of heptapeptide repeat 1 (HR1); –NH_3_ ^+^ and – COO^−^ caps are added, respectively, at *N-* and *C-*termini of each protomer.

In the case of the B.1.1.7 variant, gaps left by deletions in all three protomers are replaced with artificially long C–N bonds; systems are then allowed to relax with a 400-step preminimization cycle *in vacuo* (200 steepest-descent + 200 conjugate gradient), using the *AMBER* platform’s *sander* utility (version 18) ^[24]^, in which harmonic positional restraints (*k =* 5.0 kcal mol^−1^ Å^−2^) are applied to all atoms except those in the five residues on either side of the gap. Distortions and clashes introduced with the glycosylated Ser13–Pro26 fragment are resolved using a similar approach.

The artificial PT188-EM was modeled following the methods described in ^[5b]^.

### MD Simulation Details

After preparation, glycosylated S protein structures are solvated in a cuboidal box of TIP3P water molecules using *AMBER*’s *tleap* tool; where necessary, Na^+^ or Cl^−^ ions are added accordingly to neutralize the charge. *N-*glycosylated asparagines and oligosaccharides are treated using the *GLYCAM-06j* forcefield^[25]^, whereas ions are modeled with parameters by Joung and Cheatham^[26]^. To all other (protein) atoms, we apply the *ff14SB* forcefield ^[27]^. Starting structures and topologies for all simulated variants are electronically provided as Supporting Information.

On each glycosylated S protein variant, we conduct 4 independently replicated atomistic molecular dynamics simulations (MD), using the *AMBER* package (version 18): each replica consists of two 300-step rounds of minimization, 2.069 ns preproduction, and 1 µs production. The *sander* MD engine^[24]^ is used into the earlier stages of preproduction; thereafter, we switch to the GPU-accelerated *pmemd*.*cuda* ^[24]^.

### Details on MD production

The 1 µs production stage is carried out in the *NpT* ensemble (*T* = 300 K; *p* = 1 atm) using a 2 fs time step; a cutoff of 8.0 Å is applied for the calculation of Lennard-Jones and Coulomb interactions alike. Coulomb interactions beyond this limit are computed using the Particle Mesh Ewald method ^[28]^. All bonds containing hydrogen are restrained using the *SHAKE* algorithm ^[29]^. Constant pressure is enforced *via* Berendsen’s barostat ^[30]^ with a 1 ps relaxation time, whereas temperature is stabilized by Langevin’s thermostat^[31]^ with a 5 ps^−1^ collision frequency.

### Details on MD preproduction

Prior to the production stage, every independent MD replica for every S variant goes through a series of preproduction steps, namely: minimization, solvent equilibration, system heating, and equilibration. The first two are conducted using the *sander* utility, after which the GPU-accelerated *pmemd*.*cuda* is invoked instead.

Minimization takes place in two 300-step rounds, the first 10 of which use the steepest-descent algorithm and the last 290 conjugate gradient. In the first round, we only minimize backbone Hα and H1 hydrogens on aminoacids and monosaccharides, respectively, restraining all other atoms harmonically (*k* = 5.0 kcal mol^−1^ Å^−2^). Thereafter, all atoms are released, including solvent and ions.

Solvent equilibration occurs over 9 ps with a time step of 1 fs; the ensemble is *NVT*, with temperatures in this case enforced by the Berendsen thermostat ^[30]^. Positions of non-solvent atoms are harmonically restrained (*k* = 10 kcal mol^−1^ Å^−2^). Solvent molecules are assigned initial random velocities to match a temperature of 25 K. Fast heating to 400 K (coupling: 0.2 ps) is performed over the first 3 ps; the solvent is then retained at 400 K for another 3 ps; and cooled back down to 25 K over the last 3 ps, more slowly (coupling: 2.0). The cutoff for determining Lennard-Jones and Coulomb interactions remains at 8.0 Å for this and all subsequent stages, as does the Particle Mesh Ewald method^[28]^ to determine Coulomb interactions beyond this cutoff. *SHAKE* constraints ^[29]^ are not applied at this stage, but are always present thereafter.

For system heating, the time step is increased to 2 fs and, whilst continuing in the *NVT* ensemble, temperatures are now enforced by the Langevin thermostat^[31]^ (which remains in place for all subsequent stages). With an initial collision frequency of 0.75 ps^−1^, the system is heated from 25 to 300 K over 20 ps: all atoms are free to move except aminoacids’ Cα atoms, which are positionally restrained with *k* = 5 kcal mol^−1^ Å^−2^.

For equilibration, the ensemble is switched to *NpT* (*p* = 1 atm; Berendsen barostatcoupling: 1 ps), and the system is simulated for a further 2040 ps. The thermostat’s collision frequency is kept lower than in the production stage (1 ps^−1^). Restraints on Cα atoms are lifted gradually: *k* = 3.75 kcal mol^− 1^ Å^−2^ for the first 20 ps; 1.75 kcal mol^−1^ Å^−2^ for the following 20 ps; none thereafter.

### Clustering of MD Simulations

Following MD, each variant’s 4 replicas are concatenated into a single 4 µs ‘metatrajectory’, desolvated, stripped of any ions, and aligned on backbone heavy atoms of all aminoacid residues, in all three protomers, that belong to neither the NTD nor the RBD according to domain definitions by Huang *et al*. ^[32]^ Clustering calculations are then conducted using the hierarchical agglomerative algorithm^[33]^, considering every 20^th^ metatrajectory frame (*i*.*e*., every 50 ps), based on the root-mean-square deviation of backbone heavy atoms of aminoacid residues composing the NTD and the RBD in all three protomers. Values of ε are chosen so that they provide the best compromise between maximizing cluster homogeneity, based on silhouette score, and ensuring at least 60-80% of the metatrajectory is covered by the three most populated clusters: this usually means ε=9-12.

All of the steps discussed in the previous paragraph are conducted using *AMBER*’s postprocessing utility *cpptraj*.

### MLCE method

Potential epitopes on each S variant are predicted using the Matrix of Low Coupling Energies (MLCE) method (of which we also provide a more detailed account in our previous work) ^[15h]^. The procedure is automatically carried out by our own in-house code (https://github.com/colombolab/MLCE) which we have now rewritten to rely on the computationally more efficient MMPBSA.py utility ^[34]^ instead of mm_pbsa.pl.

To begin with, three representative S protein structures for each variant (i.e., the centroids of its three most populated clusters; see *Clustering of MD Simulations*) are minimized for 200 steps (10 steepest descent; 190 conjugate gradient), with the *sander* engine, in implicit solvent (*per* the modified Generalized Born model by Onufriev *et al*.) ^[35]^. The cutoff used for the computation of Lennard-Jones and Coulomb interactions is 12.0 Å, under nonperiodic conditions; the mobile counterion concentration is always set at 0.1 M for all variants, and the solvent-accessible surface area (SASA) is calculated by employing linear combinations of pairwise overlaps (LCPO)

After minimization, MMPBSA.py ^[34]^ is initialized and uses the MM/GBSA method ^[36]^ to construct a nonbonded pairwise interaction matrix **M**_*s*_ for each of the variant’s representative structures *s*: otherwise put, each term *M*_*s,ij*_ in this matrix contains the van der Waals and Coulomb interactions (including 1−4 interaction terms) for a representative structure’s *i*^th^ and *j*^th^ residues (aminoacids and monosaccharides alike). Settings for this stage are identical to minimization, apart from a 0 M implicit ion concentration. **M**_*s*_ matrices for a variant’s representative structures *s =* 1, 2, and 3 are then averaged, and scaled by the fraction of that variant’s trajectory represented by their parent cluster: this gives an average weighted nonbonded interaction matrix **M**.

Using the validated approach explained in detail by Genoni and coworkers ^[37]^, our code performs eigendecomposition of **M**: in other words, each averaged interaction matrix element *M*_*ij*_ —i.e., for each individual pair formed by the *i*^th^ and *j*^th^ residues—is first re-expressed as

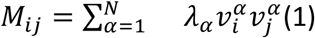

where *N* is the total number of aminoacid and glycan residues, λ_α_ is the α^th^ eigenvalue, and 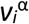 and 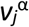 are the *i*^th^ and *j*^th^ components of its associated vector. From this eigendecomposition, the code is programmed to (re)select a minimum number of *essential eigenvectors N*_*e*_ that are sufficient to ‘cover’ interactions of the maximum number of residues in the protein. This step results in an “essential folding matrix” **M**^**fold**^ constructed from **M**’s *N*_*e*_ essential eigenvectors, and whose elements are each expressed as follows

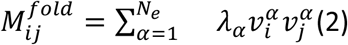

i.e., no longer as a sum over all vectors (residues) *N*, but only over the chosen *N*_*e*_ essential eigenvectors. **M**^**fold**^ thus only contains information on whether interaction of a variant’s *i*^th^ and *j*^th^’s residues is actually more or less stabilizing for the folding of one of that variant’s domains^[37]^ (e.g., NTD, RBD, etc.). This is in contrast to **M**, whose elements *M*_*ij*_ simply indicate whether a variant’s *i*^th^ and *j*^th^’s residues attract, repel, or don’t interact.

To obtain the final **MLCE** matrix, the essential folding matrix **M**^**fold**^ is subsequently Hadamard-multiplied by a pairwise residue-residue contact matrix **C**:

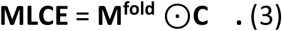

**C**’s elements *C*_*ij*_ are either 0 or 1, depending on whether Cβ atoms in the *i*^th^ and *j*^th^ residues (C1 in the case of monosaccharides; H in the case of glycines) fall below or above an arbitrary 6.0 Å threshold, respectively. In this way, each **MLCE** element *MLCE*_*ij*_ will be nonzero if and only if residues *i* and *j* are spatially contiguous *and* exhibit an energetic interaction that is stabilizing for the folding of a particular domain, in which case *MLCE*_*ij*_ takes the actual value of the pair’s degree of stabilization.

Individual *MLCE*_*ij*_ elements are ultimately ranked from the most stabilizing (i.e., pairs that are most energetically relevant for the folding of their particular domain) to least stabilizing (i.e., pairs showing the weakest energetic coupling within their domains). Our final epitope predictions are made by isolating the top 10% weakest-interacting spatially contiguous residue pairs yielded by this ranking.

## Acknowledgements

The authors thank Prof. Robert J. Woods and Dr. Oliver Grant (University of Georgia) for kindly providing atomistic molecular dynamics simulations of the fully glycosylated spike proteins. GC gratefully acknowledges Dr. Guido Scarabelli, Dr. Riccardo Capelli, and Dr. Claudio Peri for previous work on epitope prediction. This research was partially supported by a Grant from “Cassa di Risparmio di Padova e Rovigo (CARIPARO) PROGETTI DI RICERCA SUL COVID-19”.

