## Supplemental Table 1, Supplemental Table 2 for "SARS-CoV-2 Spike Protein Mutations and Escape from Antibodies: a Computational Model of Epitope Loss in Variants of Concern"

**Table S1.** The complete references to papers in which the various Abs targeting the Spike protein and its variants have been characterized and used for reference in the paper.

| Anticorpo | PDB code | D614G | B.1.1.7 | 501Y.V2.noΔ | P.1 | N439K | N501Y | PT188-EM |
|---|---|---|---|---|---|---|---|---|
| **REGN10987** | 6XDG | Hansen[1] | Tada[2] | Tada[2] | Wang[3] | Thomson[4] | Luan[5] | |
| **REGN10933** | 6XDG | Hansen[1] | Tada[2] | Tada[2] | Wang[3] | Thomson[4] | Luan[5] | |
| **LY-CoV555** | 7L3N | Jones[6] | Wang[7] | Wang[7] | | Thomson[4] | Luan[5] | |
| **S309** | 7JX3/6WPT | Pinto[8] | Wang[7] | Wang[7] | Wang[3] | Thomson[4] | Luan[5] | |
| **C135** | 7K8Z | Barnes[9] | Wang[7] | Wang[7] | | Weisblum[10] | | |
| **C144** | 7K90 | Barnes[9] | | Martinez[11] | Weisblum[10] | Weisblum[10] | | |
| **C121** | 7K90 | Barnes[9] | | Martinez[11] | Wang[3] | Weisblum[10] | | |
| **4A8** | 7C2L | Chi[12] | Wang[13] | Wang[13] | Wang[14] | | | Andreano[15] |
| **DH1041** | 7LAA | Gobeil[16] | Gobeil[16] | Gobeil[16] | Shen[17] | | | |
| **DH1043** | 7LJR | Gobeil[16] | Gobeil[16] | Gobeil[16] | Shen[17] | | | |
| **DH1047** | 7LD1 | Gobeil[16] | Gobeil[16] | Gobeil[16] | Shen[17] | | | |
| **DH1050.1** | 7LCN | Gobeil[16] | Gobeil[16] | Gobeil[16] | Shen[17] | | | |
| **S2M11** | 7K43 | Tortici[18] | | Collier[19] | Collier[19] | | | |
| **COVA1-16** | 7JMX | Wang[13] | Wang[13] | Wang[13] | Liu[20] | | Wang[13] | |
| **B38** | 7BZ5 | Wu[21] | Shen[17] | | | | Luan[5] | |
| **C002** | 7K8T | Barnes[9] | | | | | | |
| **CB6(LY-CoV016)** | 7C01 | Shi[22] | Wang[7] | Wang[7] | Wang[3] | | Luan[5] | |


1. Hansen, J.; Baum, A.; Pascal, K. E.; Russo, V.; Giordano, S.; Wloga, E.; Fulton, B. O.; Yan, Y.; Koon, K.; Patel, K.; Chung, K. M.; Hermann, A.; Ullman, E.; Cruz, J.; Rafique, A.; Huang, T.; Fairhurst, J.; Libertiny, C.; Malbec, M.; Lee, W. Y.; Welsh, R.; Farr, G.; Pennington, S.; Deshpande, D.; Cheng, J.; Watty, A.; Bouffard, P.; Babb, R.; Levenkova, N.; Chen, C.; Zhang, B.; Romero Hernandez, A.; Saotome, K.; Zhou, Y.; Franklin, M.; Sivapalasingam, S.; Lye, D. C.; Weston, S.; Logue, J.; Haupt, R.; Frieman, M.; Chen, G.; Olson, W.; Murphy, A. J.; Stahl, N.; Yancopoulos, G. D.; Kyratsous, C. A., Studies in humanized mice and convalescent humans yield a SARS-CoV-2 antibody cocktail. *Science* **2020,** *369* (6506), 1010-1014.

2. Tada, T.; Dcosta, B. M.; Zhou, H.; Vaill, A.; Kazmierski, W.; Landau, N. R., Decreased neutralization of SARS-CoV-2 global variants by therapeutic anti-spike protein monoclonal antibodies. *bioRxiv* **2021**.
3. Wang, P.; Casner, R. G.; Nair, M. S.; Wang, M.; Yu, J.; Cerutti, G.; Liu, L.; Kwong, P. D.; Huang, Y.; Shapiro, L.; Ho, D. D., Increased resistance of SARS-CoV-2 variant P.1 to antibody neutralization. *Cell Host Microbe* **2021,** *29* (5), 747-751.e4.
4. Thomson, E. C.; Rosen, L. E.; Shepherd, J. G.; Spreafico, R.; da Silva Filipe, A.; Wojcechowskyj, J. A.; Davis, C.; Piccoli, L.; Pascall, D. J.; Dillen, J.; Lytras, S.; Czudnochowski, N.; Shah, R.; Meury, M.; Jesudason, N.; De Marco, A.; Li, K.; Bassi, J.; O'Toole, A.; Pinto, D.; Colquhoun, R. M.; Culap, K.; Jackson, B.; Zatta, F.; Rambaut, A.; Jaconi, S.; Sreenu, V. B.; Nix, J.; Zhang, I.; Jarrett, R. F.; Glass, W. G.; Beltramello, M.; Nomikou, K.; Pizzuto, M.; Tong, L.; Cameroni, E.; Croll, T. I.; Johnson, N.; Di Iulio, J.; Wickenhagen, A.; Ceschi, A.; Harbison, A. M.; Mair, D.; Ferrari, P.; Smollett, K.; Sallusto, F.; Carmichael, S.; Garzoni, C.; Nichols, J.; Galli, M.; Hughes, J.; Riva, A.; Ho, A.; Schiuma, M.; Semple, M. G.; Openshaw, P. J. M.; Fadda, E.; Baillie, J. K.; Chodera, J. D.; Rihn, S. J.; Lycett, S. J.; Virgin, H. W.; Telenti, A.; Corti, D.; Robertson, D. L.; Snell, G.; Investigators, I. C.; Consortium, C.-G. U. C.-U., Circulating SARS-CoV-2 spike N439K variants maintain fitness while evading antibody-mediated immunity. *Cell* **2021,** *184* (5), 1171-1187.e20.
5. Luan, B.; Wang, H.; Huynh, T., Enhanced binding of the N501Y-mutated SARS-CoV-2 spike protein to the human ACE2 receptor: insights from molecular dynamics simulations. *FEBS Lett* **2021,** *595* (10), 1454-1461.
6. Jones, B. E.; Brown-Augsburger, P. L.; Corbett, K. S.; Westendorf, K.; Davies, J.; Cujec, T. P.; Wiethoff, C. M.; Blackbourne, J. L.; Heinz, B. A.; Foster, D.; Higgs, R. E.; Balasubramaniam, D.; Wang, L.; Bidshahri, R.; Kraft, L.; Hwang, Y.; Žentelis, S.; Jepson, K. R.; Goya, R.; Smith, M. A.; Collins, D. W.; Hinshaw, S. J.; Tycho, S. A.; Pellacani, D.; Xiang, P.; Muthuraman, K.; Sobhanifar, S.; Piper, M. H.; Triana, F. J.; Hendle, J.; Pustilnik, A.; Adams, A. C.; Berens, S. J.; Baric, R. S.; Martinez, D. R.; Cross, R. W.; Geisbert, T. W.; Borisevich, V.; Abiona, O.; Belli, H. M.; de Vries, M.; Mohamed, A.; Dittmann, M.; Samanovic, M.; Mulligan, M. J.; Goldsmith, J. A.; Hsieh, C. L.; Johnson, N. V.; Wrapp, D.; McLellan, J. S.; Barnhart, B. C.; Graham, B. S.; Mascola, J. R.; Hansen, C. L.; Falconer, E., LY-CoV555, a rapidly isolated potent neutralizing antibody, provides protection in a non-human primate model of SARS-CoV-2 infection. *bioRxiv* **2020**.
7. Wang, L.; Zhou, T.; Zhang, Y.; Yang, E. S.; Schramm, C. A.; Shi, W.; Pegu, A.; Oloniniyi, O. K.; Henry, A. R.; Darko, S.; Narpala, S. R.; Hatcher, C.; Martinez, D. R.; Tsybovsky, Y.; Phung, E.; Abiona, O. M.; Antia, A.; Cale, E. M.; Chang, L. A.; Choe, M.; Corbett, K. S.; Davis, R. L.; DiPiazza, A. T.; Gordon, I. J.; Helmold Hait, S.; Hermanus, T.; Kgagudi, P.; Laboune, F.; Leung, K.; Liu, T.; Mason, R. D.; Nazzari, A. F.; Novik, L.; O'Connell, S.; O'Dell, S.; Olia, A. S.; Schmidt, S. D.; Stephens, T.; Stringham, C. D.; Talana, C. A.; Teng, I. T.; Wagner, D. A.; Widge, A. T.; Zhang, B.; Roederer, M.; Ledgerwood, J. E.; Ruckwardt, T. J.; Gaudinski, M. R.; Moore, P. L.; Doria-Rose, N. A.; Baric, R. S.; Graham, B. S.; McDermott, A. B.; Douek, D. C.; Kwong, P. D.; Mascola, J. R.; Sullivan, N. J.; Misasi, J., Ultrapotent antibodies against diverse and highly transmissible SARS-CoV-2 variants. *Science* **2021**.
8. Pinto, D.; Park, Y. J.; Beltramello, M.; Walls, A. C.; Tortorici, M. A.; Bianchi, S.; Jaconi, S.; Culap, K.; Zatta, F.; De Marco, A.; Peter, A.; Guarino, B.; Spreafico, R.; Cameroni, E.; Case, J. B.; Chen, R. E.; Havenar-Daughton, C.; Snell, G.; Telenti, A.; Virgin, H. W.; Lanzavecchia, A.; Diamond, M. S.; Fink, K.; Veesler, D.; Corti, D., Cross-neutralization of SARS-CoV-2 by a human monoclonal SARS-CoV antibody. *Nature* **2020,** *583* (7815), 290-295.
9. Barnes, C. O.; Jette, C. A.; Abernathy, M. E.; Dam, K. A.; Esswein, S. R.; Gristick, H. B.; Malyutin, A. G.; Sharaf, N. G.; Huey-Tubman, K. E.; Lee, Y. E.; Robbiani, D. F.; Nussenzweig, M.

C.; West, A. P.; Bjorkman, P. J., SARS-CoV-2 neutralizing antibody structures inform therapeutic strategies. *Nature* **2020,** *588* (7839), 682-687.
10. Weisblum, Y.; Schmidt, F.; Zhang, F.; DaSilva, J.; Poston, D.; Lorenzi, J. C.; Muecksch, F.; Rutkowska, M.; Hoffmann, H. H.; Michailidis, E.; Gaebler, C.; Agudelo, M.; Cho, A.; Wang, Z.; Gazumyan, A.; Cipolla, M.; Luchsinger, L.; Hillyer, C. D.; Caskey, M.; Robbiani, D. F.; Rice, C. M.; Nussenzweig, M. C.; Hatziioannou, T.; Bieniasz, P. D., Escape from neutralizing antibodies by SARS-CoV-2 spike protein variants. *Elife* **2020,** *9*.
11. Martinez, D. R.; Schaefer, A.; Leist, S. R.; Gully, K.; Feng, J. Y.; Bunyan, E.; Porter, D. P.; Cihlar, T.; Montgomery, S. A.; Baric, R. S.; Nussenzweig, M. C.; Sheahan, T. P., Early therapy with remdesivir and antibody combinations improves COVID-19 disease in mice. *bioRxiv* **2021**.
12. Chi, X.; Yan, R.; Zhang, J.; Zhang, G.; Zhang, Y.; Hao, M.; Zhang, Z.; Fan, P.; Dong, Y.; Yang, Y.; Chen, Z.; Guo, Y.; Li, Y.; Song, X.; Chen, Y.; Xia, L.; Fu, L.; Hou, L.; Xu, J.; Yu, C.; Li, J.; Zhou, Q.; Chen, W., A neutralizing human antibody binds to the N-terminal domain of the Spike protein of SARS-CoV-2. *Science* **2020,** *369* (6504), 650-655.
13. Wang, P.; Nair, M. S.; Liu, L.; Iketani, S.; Luo, Y.; Guo, Y.; Wang, M.; Yu, J.; Zhang, B.; Kwong, P. D.; Graham, B. S.; Mascola, J. R.; Chang, J. Y.; Yin, M. T.; Sobieszczyk, M.; Kyratsous, C. A.; Shapiro, L.; Sheng, Z.; Huang, Y.; Ho, D. D., Antibody resistance of SARS-CoV-2 variants B.1.351 and B.1.1.7. *Nature* **2021,** *593* (7857), 130-135.
14. Wang, R.; Zhang, Q.; Ge, J.; Ren, W.; Zhang, R.; Lan, J.; Ju, B.; Su, B.; Yu, F.; Chen, P.; Liao, H.; Feng, Y.; Li, X.; Shi, X.; Zhang, Z.; Zhang, F.; Ding, Q.; Zhang, T.; Wang, X.; Zhang, L., Analysis of SARS-CoV-2 variant mutations reveals neutralization escape mechanisms and the ability to use ACE2 receptors from additional species. *Immunity* **2021**.
15. Andreano, E.; Nicastri, E.; Paciello, I.; Pileri, P.; Manganaro, N.; Piccini, G.; Manenti, A.; Pantano, E.; Kabanova, A.; Troisi, M.; Vacca, F.; Cardamone, D.; De Santi, C.; Torres, J. L.; Ozorowski, G.; Benincasa, L.; Jang, H.; Di Genova, C.; Depau, L.; Brunetti, J.; Agrati, C.; Capobianchi, M. R.; Castilletti, C.; Emiliozzi, A.; Fabbiani, M.; Montagnani, F.; Bracci, L.; Sautto, G.; Ross, T. M.; Montomoli, E.; Temperton, N.; Ward, A. B.; Sala, C.; Ippolito, G.; Rappuoli, R., Extremely potent human monoclonal antibodies from COVID-19 convalescent patients. *Cell* **2021,** *184* (7), 1821-1835.e16.
16. Gobeil, S. M.; Janowska, K.; McDowell, S.; Mansouri, K.; Parks, R.; Stalls, V.; Kopp, M. F.; Manne, K.; Li, D.; Wiehe, K.; Saunders, K. O.; Edwards, R. J.; Korber, B.; Haynes, B. F.; Henderson, R.; Acharya, P., Effect of natural mutations of SARS-CoV-2 on spike structure, conformation, and antigenicity. *Science* **2021**.
17. Shen, X.; Tang, H.; McDanal, C.; Wagh, K.; Fischer, W.; Theiler, J.; Yoon, H.; Li, D.; Haynes, B. F.; Sanders, K. O.; Gnanakaran, S.; Hengartner, N.; Pajon, R.; Smith, G.; Glenn, G. M.; Korber, B.; Montefiori, D. C., SARS-CoV-2 variant B.1.1.7 is susceptible to neutralizing antibodies elicited by ancestral spike vaccines. *Cell Host Microbe* **2021,** *29* (4), 529-539.e3.
18. Tortorici, M. A.; Beltramello, M.; Lempp, F. A.; Pinto, D.; Dang, H. V.; Rosen, L. E.; McCallum, M.; Bowen, J.; Minola, A.; Jaconi, S.; Zatta, F.; De Marco, A.; Guarino, B.; Bianchi, S.; Lauron, E. J.; Tucker, H.; Zhou, J.; Peter, A.; Havenar-Daughton, C.; Wojcechowskyj, J. A.; Case, J. B.; Chen, R. E.; Kaiser, H.; Montiel-Ruiz, M.; Meury, M.; Czudnochowski, N.; Spreafico, R.; Dillen, J.; Ng, C.; Sprugasci, N.; Culap, K.; Benigni, F.; Abdelnabi, R.; Foo, S. C.; Schmid, M. A.; Cameroni, E.; Riva, A.; Gabrieli, A.; Galli, M.; Pizzuto, M. S.; Neyts, J.; Diamond, M. S.; Virgin, H. W.; Snell, G.; Corti, D.; Fink, K.; Veesler, D., Ultrapotent human antibodies protect against SARS-CoV-2 challenge via multiple mechanisms. *Science* **2020,** *370* (6519), 950-957.
19. Collier, D. A.; De Marco, A.; Ferreira, I. A. T. M.; Meng, B.; Datir, R. P.; Walls, A. C.; Kemp, S. A.; Bassi, J.; Pinto, D.; Silacci-Fregni, C.; Bianchi, S.; Tortorici, M. A.; Bowen, J.; Culap, K.; Jaconi, S.; Cameroni, E.; Snell, G.; Pizzuto, M. S.; Pellanda, A. F.; Garzoni, C.; Riva, A.; Elmer, A.;

Kingston, N.; Graves, B.; McCoy, L. E.; Smith, K. G. C.; Bradley, J. R.; Temperton, N.; Ceron-Gutierrez, L.; Barcenas-Morales, G.; Harvey, W.; Virgin, H. W.; Lanzavecchia, A.; Piccoli, L.; Doffinger, R.; Wills, M.; Veesler, D.; Corti, D.; Gupta, R. K.; Collaboration, C.-N. B. C.-.; Consortium, C.-G. U. C.-U., Sensitivity of SARS-CoV-2 B.1.1.7 to mRNA vaccine-elicited antibodies. *Nature* **2021,** *593* (7857), 136-141.
20. Liu, H.; Wu, N. C.; Yuan, M.; Bangaru, S.; Torres, J. L.; Caniels, T. G.; van Schooten, J.; Zhu, X.; Lee, C. D.; Brouwer, P. J. M.; van Gils, M. J.; Sanders, R. W.; Ward, A. B.; Wilson, I. A., Cross-Neutralization of a SARS-CoV-2 Antibody to a Functionally Conserved Site Is Mediated by Avidity. *Immunity* **2020,** *53* (6), 1272-1280.e5.
21. Wu, Y.; Wang, F.; Shen, C.; Peng, W.; Li, D.; Zhao, C.; Li, Z.; Li, S.; Bi, Y.; Yang, Y.; Gong, Y.; Xiao, H.; Fan, Z.; Tan, S.; Wu, G.; Tan, W.; Lu, X.; Fan, C.; Wang, Q.; Liu, Y.; Zhang, C.; Qi, J.; Gao, G. F.; Gao, F.; Liu, L., A noncompeting pair of human neutralizing antibodies block COVID-19 virus binding to its receptor ACE2. *Science* **2020,** *368* (6496), 1274-1278.
22. Shi, R.; Shan, C.; Duan, X.; Chen, Z.; Liu, P.; Song, J.; Song, T.; Bi, X.; Han, C.; Wu, L.; Gao, G.; Hu, X.; Zhang, Y.; Tong, Z.; Huang, W.; Liu, W. J.; Wu, G.; Zhang, B.; Wang, L.; Qi, J.; Feng, H.; Wang, F. S.; Wang, Q.; Gao, G. F.; Yuan, Z.; Yan, J., A human neutralizing antibody targets the receptor-binding site of SARS-CoV-2. *Nature* **2020,** *584* (7819), 120-124.

**Table S2.** The residues that are part of the experimentally determined epitopes and of the predicted epitopes on the various Spike variants. The numbering of the residues refers to that reported in UNIPROT entry P0DTC2. https://www.uniprot.org/uniprot/P0DTC2.

| Antibody | Experimental | D614G | N439K | 501Y.V2.noΔ | B.1.1.7 | P.1 | N501Y | PT188-EM |
|---|---|---|---|---|---|---|---|---|
| **4A8** | 143-148,150,152,245-250,256-257 | 143-147,245-246,249 | 144-147,150,152,257 | no | 143,145,147-148,245 | 143,145-148,245,248,250,256 | no | 147,150,152 |
| **DH1050.1** | 140-146,148-152,156-159,244-245 | 140,143-146,156,158-159,245 | 140-141,144-146,149-152,159,244 | no | 142-143,145,148-149,151,157,244-245 | 142-143,145-146,148-149,156-157,244-245 | no | 149-152 |
| **REGN10987** | 345-346,439-441,443-447,449,490-500 | 439,441,443-447,449,491-498 | no | 345-346,439,441,443,490-491,494-497 | 439-441,443-447,449,490,492,493-500 | 491 | no | 439-440 |
| **REGN10933** | 403,405,417,421,453,455-456,473,475-478,484-490,492-494,498 | 403,405,421,453,456,473,476,478,485,487,489,492-494,498 | 484,486-487 | 403,405,453,477,484-485,489-490,494 | 403,421,453,456,473,475,477-478,485-486,488-490,492-494,498 | 421 | 453,473,475-478,487 | 403,406 |
| **S309** | 333-341,343-345,354,356-361,441,444,509 | 334-341,354,356-361,441,444,509 | 337,340 | 333-335,337-338,340,344-345,354,358,360,441,509 | 334-338,340,354,356-361,441,509 | 335 | 336,354,356,358-359,361,509 | 357,359-360,509 |
| **LY-CoV555** | 351,449-450,452-453,456,470,472,478,481-490,492-496 | 351,449-450,452-453,456,472,478,485,487,489,492-496 | 482-484,486-487 | 351,452-453,470,472,483-485,489-490,494-496 | 351,449,452-453,456,470,472,478,482,485-486,488-490,492-496 | no | 453,470,472,478,482-483,487 | no |
| **C135** | 341-346,438-442,490 | 341,438-439,441-442 | no | 344-346,439,441-442,490 | 448-442,490 | 438 | 438 | 439-440 |
| **C144** | 342-343,367,370-374,417,436,440,449-450,452,455-456,472,483-487,498 | 367,371,373-374,436,449-450,456,472,485,487,498 | 483-484,486-487 | 367,371-374,436,472,483-485 | 372-374,449,455-456,472,485-486,498 | 367,370,373-374 | 436,472,483,487 | 436 |
| **C121** | 444,449,483-490,492-496 | 444,449,485,487,489,492-496 | 483-484,486-487 | 483-485,489-490,494-496 | 449,485-486,488-490,492-496 | no | 483,487 | no |
| **DH1041** | 351,446-452,470-472,480-486,490-494,496 | 351,446-452,471-472,480,485,491-494,496 | 480,482-484,486 | 351,448,451-452,470,472,483-485,490-491,494,496 | 351,446-449,451-452,470-472,482,485-486,490,492-494,496 | 491 | 470-472,480,482-483 | no |
| **DH1043** | 351,446-447,449-450,452,456,470,472,478-494 | 351,446-447,449-450,452,456,472,478,480,485,487,489,491-494 | 479-480,482-484,486-487 | 351,452,470,472,483-485,489-491,494 | 351,446-447,449,452,456,470,472,478-479,482,485-486,488-490,492-494 | 491 | 470,472,478-480,482-483,487 | no |
| **DH1047** | 369-372,374-380,383-384,404-405,407-409,414-416,435,499,501-505 | 369,371,374-380,383-384,404-405,407-409,414-416,435,503,505 | no | 369,371-372,374-379,383-384,404,407,409,414-416,435 | 369,372,374-379,383,405,407-408,414-416,435,499,501-503,505 | 370,374,377,380,404,416 | 375,384,416 | 380,404-405,407-409,435,501,503-505 |
| **S2M11** | 342-343,367-368,371-373,436,440-441,446-447,449,452,455-456,484-487,489-490,492-496,498 | 367,371,373-374,436,441,446-447,449,452,456,485,487,489,492-496,498 | 484,486-487 | 367-368,371-374,436,441,452,484-485,489-490,494-496 | 368,372-374,440-441,446-447,449,452,455-456,485-486,489-490,492-496,498 | 367-368,373-374 | 368,436,487 | 436,440 |
| **COVA1-16** | 368-369,371,374-385,408-409,412-416,427-429 | 368-369,371,374-381,384,408-409,412,414-416,428 | no | 368-369,371,374-379,382-385,409,412-416,427-429 | 368-369,374-379,381-383,385,408,414-416 | 368,374,377,380-382,412,416 | 368,375,384,416 | 380-381,408-409,429 |
| **B38** | 403,405-406,409,415-417,420-421,449,453,455-460,473-477,484,486-498,500-503,505 | 403,405-406,409,415-416,420-421,449,453,456,473-474,476,487,489,491-498,503,505 | 484,486-487 | 403,406,409,420,453,474,477,484,489-491,494-497 | 403,405,415,421,449,453,455,457-460,473-476,484,487-480,501-498,500-502,505 | 416,420-421,460,491 | 416,453,473-477,487 | 403,405-406,409,501,505 |
| **C002** | 444,447,449-450,452,470,473,481-490,492-494 | 444,447,449-450,452,473,485,487,489,492-494 | 482-484,486-487 | 452,470,483-485,489-490,494 | 447,449,452,470,473,482,485-486,488-490,492-495 | no | 470,473,482-483,487 | no |
| **CB6 (LY-CoV016)** | 403,405-406,408-409,415-417,420-421,453,455-460,473-477,486-487,489,493-495,500-505 | 403,405-406,408-409,415-416,420-421,453,456,473-474,476,487,489,493-495,503,505 | 486-487 | 403,406,409,415-416,420,453,474,477,489,494-495 | 403,405,408,415,421,453,455-459,473-475,478,486,489,493-495,500-503,505 | 416,420-421,460 | 416,453,473-477,487 | 403,405-406,408-409,501,503-505 |
